# Spatial heterogeneity in myelin sheathing impacts signaling reliability and susceptibility to injury

**DOI:** 10.1101/2023.12.04.570018

**Authors:** Afroditi Talidou, Jérémie Lefebvre

**Affiliations:** Department of Biology, University of Ottawa, Ottawa, ON K1N 6N5, Canada; Krembil Research Institute, University Health Network, Toronto, ON M5T 0S8, Canada; Department of Mathematics, University of Toronto, Toronto, ON M5S 2E4, Canada

## Abstract

Axons in the mammalian brain show significant diversity in myelination motifs, displaying spatial heterogeneity in sheathing along individual axons and across brain regions. However, its impact on neural signaling and susceptibility to injury remains poorly understood. To address this, we leveraged cable theory and developed model axons replicating the myelin sheath distributions observed experimentally in different regions of the mouse central nervous system. We examined how the spatial arrangement of myelin affects propagation and predisposition to conduction failure in axons with cortical versus callosal myelination motifs. Our results indicate that regional differences in myelination significantly influence conduction timing and signaling reliability. Sensitivity of action potential propagation to the specific positioning, lengths, and ordering of myelinated and exposed segments reveals non-linear and path-dependent conduction. Furthermore, myelination motifs impact signaling vulnerability to demyelination, with callosal motifs being particularly sensitive to myelin changes. These findings highlight the crucial role of myelinating glia in brain function and disease.

## Introduction

Myelinated axons play a pivotal role in supporting and facilitating the faithful conduction of action potentials (APs). Myelin-forming oligodendrocytes (OLs) engage in the synthesis of cellular membranes enwrapping axons through a multilamellar spiral configuration (***Bradl and Lassmann, 2010; Simons and Nave, 2015***), serving as an effective insulator for the axons and enabling saltatory conduction. Individual OLs have the capacity to generate multiple myelin sheaths, with each sheath potentially varying in length and enwrapping different axons (***Chong et al., 2012; Tomassy et al., 2014***). In contrast to the conventional depiction of axons, where myelin sheaths are uniformly distributed along their length, numerous investigations have shown that axonal myelin sheathing is, in fact, highly heterogeneous. Even when controlling for variations in length and overall myelin coverage (that is, the percentage of the axon covered by myelin, referred thereafter as PMC), internodes’ number, thickness, length and spatial arrangement do vary significantly between and along given axons (***Tomassy et al., 2014; Micheva et al., 2016; Hill et al., 2018; Call and Bergles, 2021; Auer et al., 2018; Stedehouder et al., 2019; Bakiri et al., 2011; Sturrock, 1980; Ford et al., 2015; Benamer et al., 2020***). Such diversity extends within and across brain regions. For instance, as axons traverse the corpus callosum, internode lengths exhibit greater variability, whereas, in the cerebral cortex, they tend to be more uniform (***Chong et al., 2012***). Such striking differences are of prime functional significance (***Tomassy et al., 2014***), as they suggest regional differences in myelination strategies to support neural signaling. Indeed, the spatial heterogeneity resulting from the particular positions and lengths of myelin sheaths, the molecular constitution of nodes, along with variations in the distribution and density of ion and/or leak channels, collectively influence axonal conduction and the reliable propagation of APs. Such microstructural changes notably tune axonal conduction delays – the temporal interval required for an AP originating at the axon hillock to reach post-synaptic targets – which play an important role in neural circuits operation (***Fields, 2008; Talidou et al., 2022***) and to maintain brain function (***Steadman et al., 2020; Pan et al., 2020; Bengtsson et al., 2005; Xin and Chan, 2020; McKenzie et al., 2014; Mabbott et al., 2006; Nagy et al., 2004; Pujol et al., 2006***).

A crucial question that remains unanswered is the impact of such heterogeneity on the conduction of APs. Does spatial variation in myelin sheathing pattern affect conduction delays? If so, how much? Are these patterns equally resilient to variations in myelination, or are certain motifs more fragile and prone to conduction failure with changes in myelin coverage? Understanding the implications of regional myelination motifs diversity on axonal conduction would represent an important advancement in our understanding of the tactics employed by OLs to optimize neural trafficking, and in identifying which axons or part of these axons are more vulnerable to injury. An extensive corpus of scientific literature has developed mathematical models to investigate the propagation of APs along myelinated axons (***Fitzhugh, 1962; Goldman and Albus, 1968; Basser, 1993; Nygren and Halter, 1999; McIntyre et al., 2002; Gow and Devaux, 2008; Schmidt and Knösche, 2019; Naud and Longtin, 2019; Scurfield and Latimer, 2018; Babbs and Shi, 2013; Ashida and Nogueira, 2018; Grindrod and Sleeman, 1985***). Prevailing models oftentimes presuppose uniform, homogeneous myelin sheathing interspersed with exceedingly brief nodes of Ranvier. However, this convenient approximation severely limits the characterization of the effect of spatial heterogeneity in myelin sheathing along axons (***Chong et al., 2012; Tomassy et al., 2014***).

To circumvent this limitation, we developed model axons endowed with myelin sheath patterns closely matching those measured from the cerebral cortex and corpus callosum of mice (***Chong et al., 2012***). In vivo studies conducted in these regions (and others) by ***Chong et al. (2012***) revealed considerable variability in the number, length and arrangement of myelin sheaths formed by individual OLs, representing ideal starting points to ground our explorations. We hence used these data to constrain our model axons, and compared how these distinct myelination motifs shape AP conduction reliability, as well as vulnerability to failure during demyelination. Our approach integrates variability in the lengths and number of myelin sheaths and exposed segments while keeping other parameters (e.g., axonal length, radius, g-ratio, biophysical parameters, etc) constant. This was done to ensure that any observed differences in axonal conduction can be attributed exclusively to myelin placement along the axons. We leveraged and adapted cable theory, amenable to the representation of AP propagation across axons composed of myelin sheaths and exposed segments of arbitrary (and different) lengths. Specifically, we adopted a phenomenological model to simulate the rapid depolarization (spike) of the membrane potential along the exposed segments of the axon (***Ashida and Nogueira, 2018***), while we incorporated diffusion accounting for the fast transmission of APs within myelin sheaths. We further quantified how spatial heterogeneity in myelin sheathing influences the occurrence of AP conduction failures and susceptibility to conduction block.

## Results

### Modelling axons with spatially heterogeneous myelin sheathing

To characterize changes in AP conduction resulting from realistic and physiologically observed myelination motifs, we constructed nerve axon models incorporating myelin sheath heterogeneity that mirrors experimental observations (***Chong et al., 2012***) (see Methods). We sampled myelin sheath lengths matching those documented in the cerebral cortex and corpus callosum of mice (***Chong et al., 2012***) and used probabilistic tools to implement a framework that incrementally allocates these sampled sheaths to an initially fully unmyelinated (that is, bare) axon of a predetermined length (see Methods). A schematic illustrating differences between cortical and callosal myelination motifs is shown in Fig. 1A-B.

**Figure 1.**
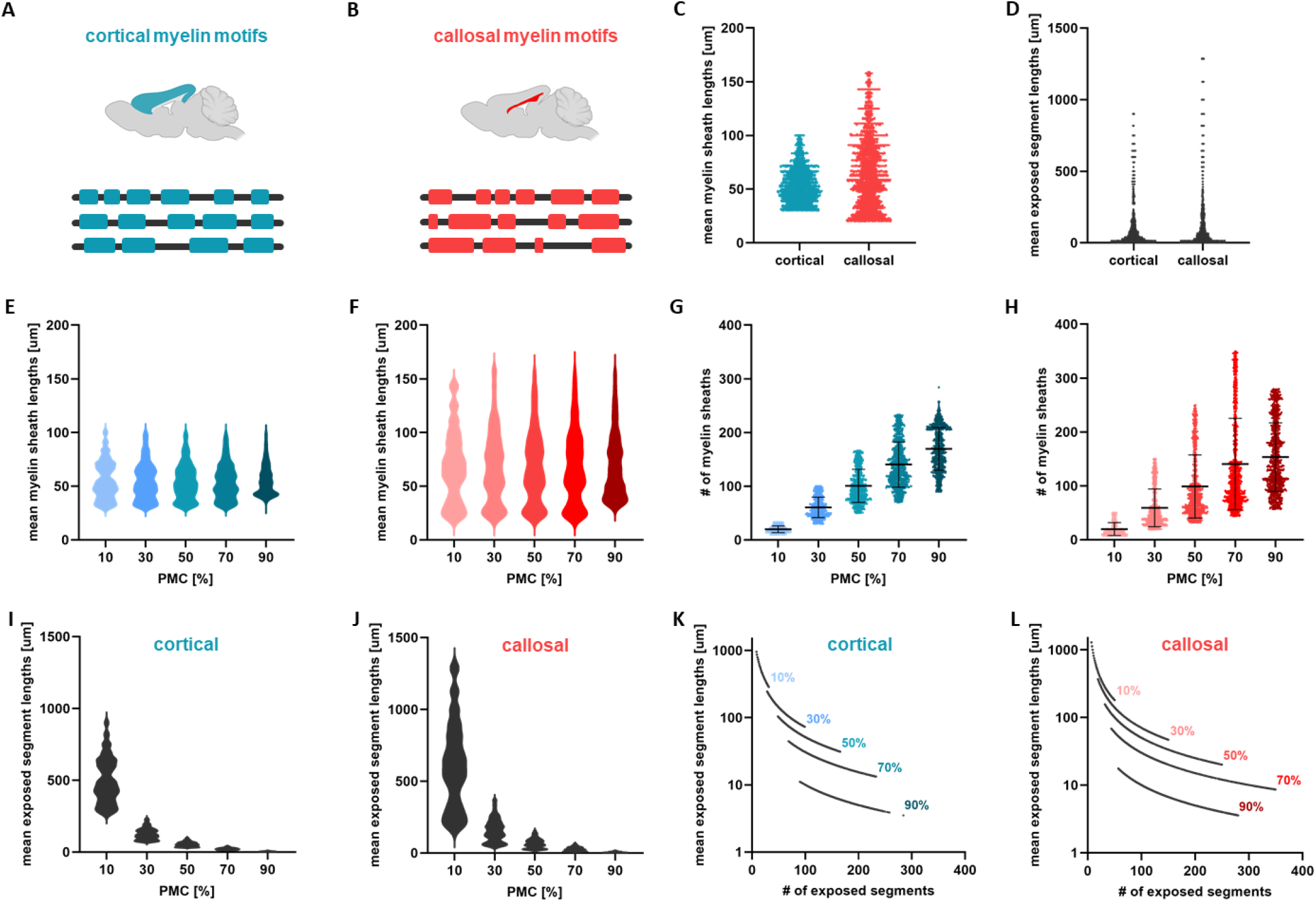
Modelling axons of heterogeneous myelination motifs in grey and white matter. **A-B**. Schematic illustration of myelination motifs along individual axons in two different brain regions: in the cerebral cortex (blue, panel **A**) and corpus callosum (red, panel **B**). Callosal motifs are more spatially heterogeneous compared to those found in the cortex. **C**. Scatter plot of the mean values of myelin sheath lengths for axons featuring cortical and callosal myelin motifs for all cases of PMC examined. **D**. Scatter plot of the mean values of exposed segment lengths. **E-F**. Mean values of myelin sheath lengths, corresponding to cortical (panel **E**) and callosal (panel **F**) motifs, plotted separately for each PMC in the form of violin plots. **G-H**. Scatter plots of the number of myelin sheaths as a function of PMC (cortical motifs - panel **G**, callosal motifs - panel **H**). The lines indicate mean values and standard deviations of the plotted data. **I-J**. Violin plots of the mean values of exposed segment lengths for each of the five cases of PMC. **K-L**. Non-linear relation between the mean values of exposed segment lengths and their number. The geometrical parameters of the axons used are: axon length = 10*mm*, radius = 2*μm* and g-ratio = 0.6. The biophysical parameter values used are given in Table 1. The results shown are for *K* = 1000 independent axons.

To broaden the applicability of our findings to axons with different levels of myelination, we examined five distinct scenarios, each representing varying PMC for a fixed total axonal length (that is, PMC values of 10%, 30%, 50%, 70% and 90%). Figure 1C-D shows the mean lengths of myelin sheaths and exposed segments for cortical and callosal myelination motifs across all PMC scenarios considered. These statistics showcase important differences between cortical and callosal myelin sheathing: myelin sheaths in the callosum display much greater spatial heterogeneity compared to those in the cortex. Specifically, as axons traverse the cortex, myelin sheaths maintain more consistent lengths. However, in the corpus callosum, the same axons display more variable myelin sheath lengths, characterized by heavier-tailed distributions (Fig. 1C), aligning with experimental measurements (***Chong et al., 2012***). In (***Chong et al., 2012; Call and Bergles, 2021***), it has been observed that the average lengths of myelin sheaths fall within a specific range, suggesting that differences in PMC primarily reflect variations in the number of sheaths per axon. Building upon this observation, we maintained the average length of myelin sheaths within a given range, specific to either cortical or callosal axons, while modifying their number. The resulting mean values of sheath lengths for each PMC value are illustrated in Fig. 1E for cortical and Fig. 1F for callosal motifs. The corresponding number of myelin sheaths is depicted in Fig. 1G-H. These clearly highlight important differences in spatial heterogeneity in both myelin sheath lengths and number between cortical and callosal motifs.

**Table 1.**
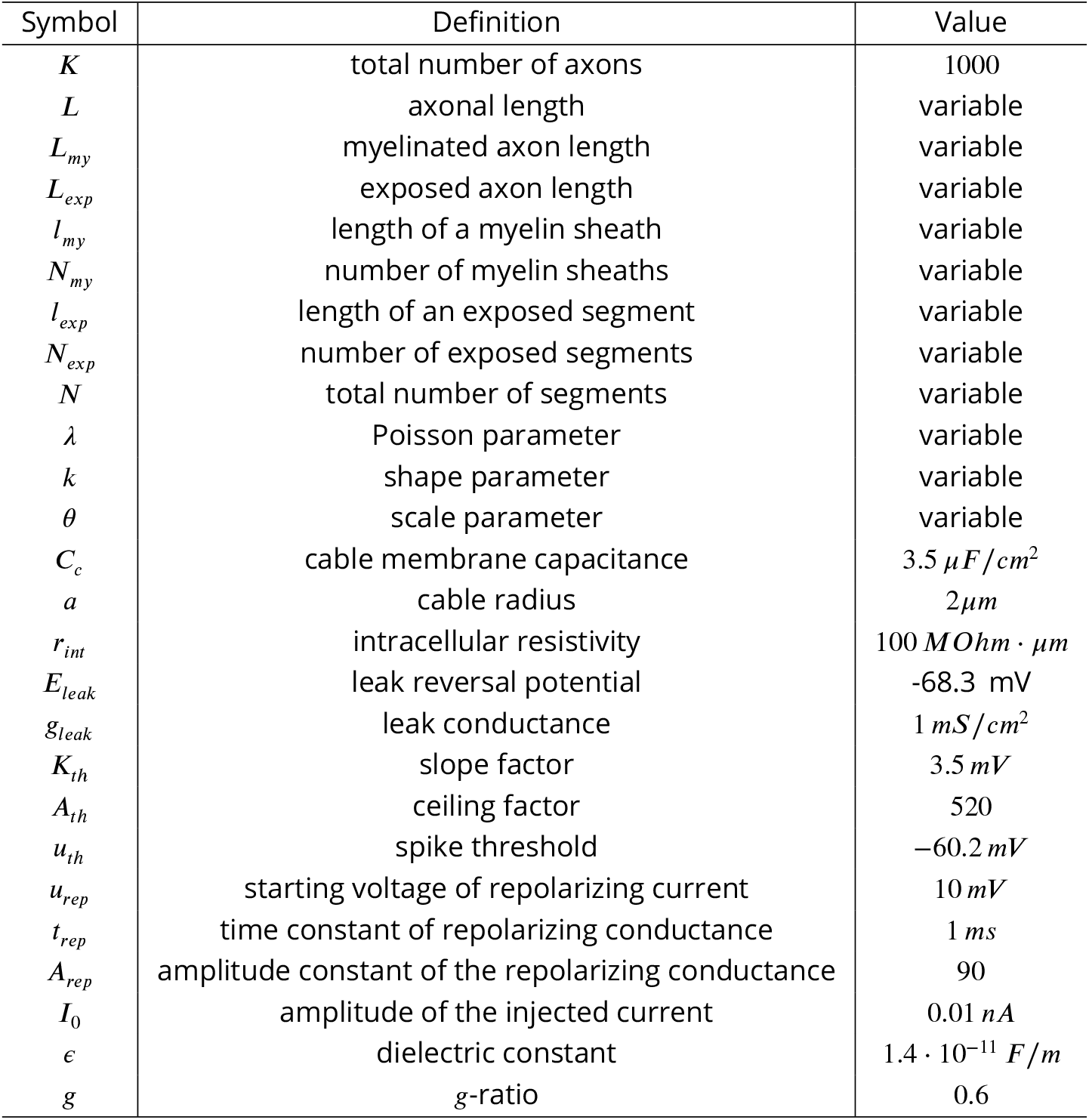
List of model variables and parameters.

Geometrically, for a given axon length, any adjustments made to the number or lengths of myelin sheaths inevitably impact the corresponding number and lengths of exposed segments. The mean lengths of exposed segments are illustrated in Fig. 1I-J. For the same value of PMC, callosal motifs exhibit significantly more variable and long exposed segments compared to cortical motifs (Fig. 1J). One may further notice that, as the PMC increases, the lengths of exposed segments decrease: as myelin covers an increasingly large portion of the axon, exposed segments become smaller and less variable. The relationship between the mean length and number of exposed segments is plotted in Fig. 1K-L. There, one can see that cortical myelination motifs (Fig. 1K) are characterized by a smaller number of exposed segments compared to callosal motifs (Fig. 1L).

To isolate the impact (if any) of myelin sheath distribution heterogeneity on AP propagation and facilitate a meaningful comparison within and between cortical and callosal motifs, we deliberately kept constant the values of axonal length, radius/caliber and myelin sheath thickness (that is, g-ratio). The biophysical parameters used (see Table 1) also remain constant. This approach introduces variability solely in the length and placement of myelin sheaths and exposed segments along the axons while ensuring that any observed differences in axonal conduction can be attributed exclusively to myelin heterogeneity.

We modelled the propagation of APs along these axons using cable theory (***Dayan and Abbott, 2005***), adapted to account for spatially heterogeneous myelination. To simulate spike generation and conduction within exposed segments, we used a spike-conducting integrate-and-fire model (***Ashida and Nogueira, 2018***). The rapid spread of APs along myelin sheaths was replicated assuming pure diffusion. The resulting system of equations is further supplemented by additional conditions that account for the current flow at the paranodal junctions, where the membrane potential transitions from myelinated to exposed segments, and vice versa (see Methods).

### Influence of myelin sheaths heterogeneity on AP conduction

We started our analysis by exploring whether myelin sheathing heterogeneity influences AP propagation along individual axons. We first compared the conduction of APs along axons displaying a homogeneous myelin motif – characterized by myelin sheaths of uniform length periodically distributed along axons – with the conduction along axons featuring spatially heterogeneous myelin sheath distributions. For the purpose of this experiment, axons exhibiting callosal myelination motifs were used. Snapshots capturing the membrane potential’s evolution over time, accompanied by schematic representations of the axons, are illustrated in Fig. 2A-D. In the homogeneous case (Fig. 2A), we observe a consistent AP waveform, displaying no discernible changes in either width or amplitude as it traverses along the axon. However, in the case of heterogeneous myelination (Fig. 2B-D), fluctuations in both the shape of the AP waveform and/or conduction time arise. These findings indeed revealed that the particular location of myelin sheaths and exposed segments as well as their number lead to instances of faster or slower conduction, or even conduction failure (Fig. 2D). Figures 2E-H present bar plots detailing the lengths of myelinated and exposed segments for each of the examined axons. For the axon with homogeneous myelination, both the lengths of myelin sheaths and exposed segments remain constant. Conversely, for the axons exhibiting heterogeneous myelination, one can see an important dispersion in those lengths.

**Figure 2.**
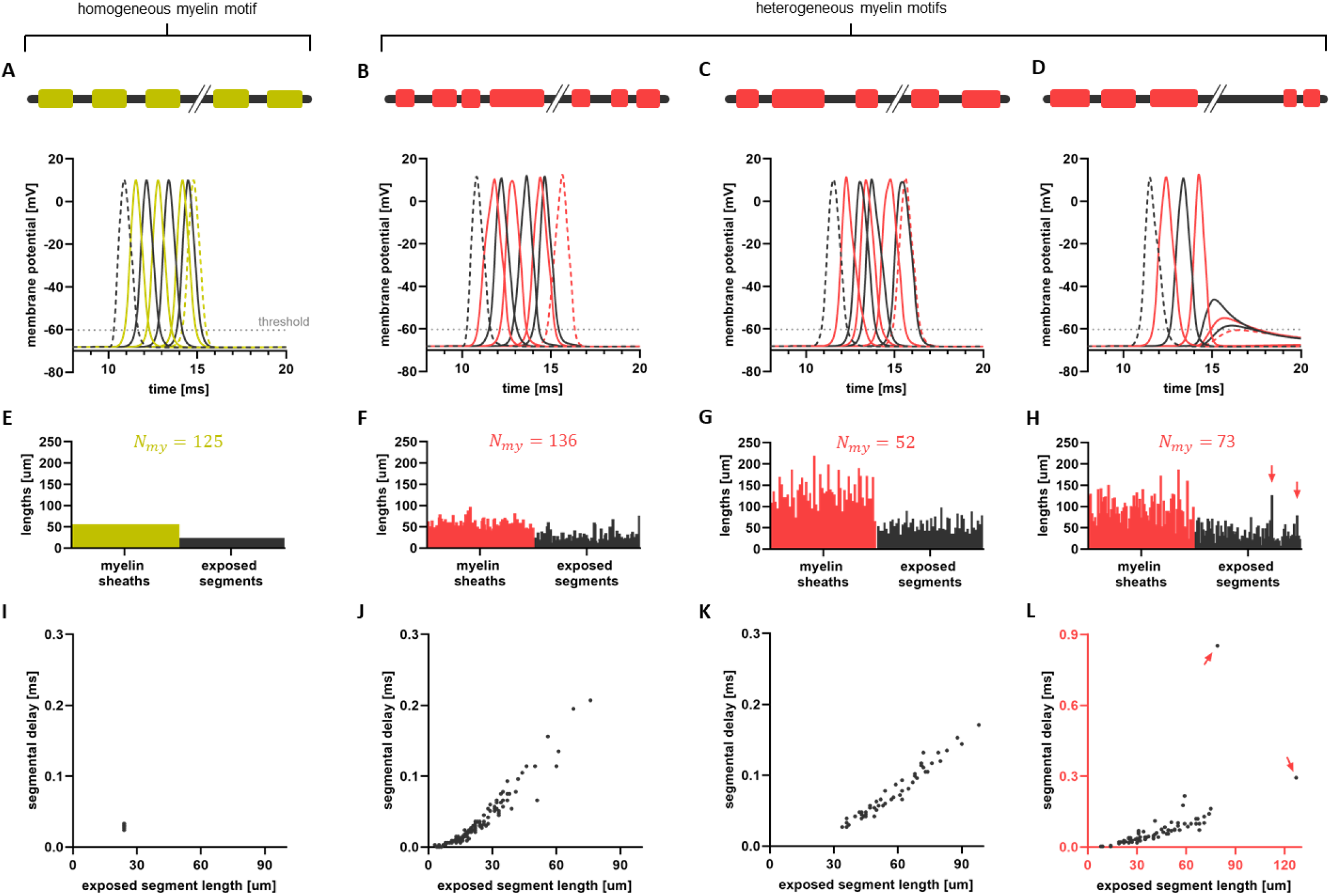
Exemplar AP propagation along axons with spatially homogeneous and heterogeneous myelin sheathing. **A**. Snapshots illustrating the propagation of AP along an axon characterized by homogeneous myelination. **B-D**. Propagation of APs along axons exhibiting heterogeneous myelin motifs. For all four examples, the axonal length is *L* = 10 *mm* and PMC is 70 %. A schematic representation of each axon is displayed at the top of these panels. **E-H**. Bar charts of the lengths of myelin sheaths and exposed segments for the four individual axons examined. Each bar in panels **F-H** represents the length of either a myelin sheath (red) or an exposed segment (black). As there is no variability in the lengths of sheaths and exposed segments along axons of homogeneous myelination, all those values are constant (panel **E**). The number of myelin sheaths varies for each axon and is denoted by *N*_*my*_. The number of exposed segments (*N*_*exp*_) equals the number of myelin sheaths. In panel **E**, where the length of myelin sheaths is 56 *μm*, resulting in a total of 125 sheaths, assuming a uniform distribution of exposed segments, their individual length is 24 *μm*. Note, however, that the number and length of myelin sheaths and exposed segments vary across all three axons exhibiting heterogeneous myelin motifs. **I-L**. The delays of APs along individual exposed segments are plotted as a function of the length of those segments. In the homogeneous case, these values remain constant, as depicted in panel **I**, whereas in the heterogeneous cases, they exhibit great variability. In panels **H** and **L**, the red arrows highlight the largest exposed segments, which result in significant delays and ultimately lead to conduction block.

These observations suggest that variability in the lengths and number of myelin sheaths and exposed segments can result in fluctuations in both the waveform and propagation of APs, thereby also influencing the axonal conduction delay. Indeed, segmental delays – representing the time required for an AP to traverse a myelinated and/or exposed segment (see Methods) – were found to vary with the length of each segment in heterogeneous motifs, as depicted in Fig. 2J-L. Note that the slope in each of the panels of Fig. 2J-L is different reflecting to fluctuations in conduction velocities (CVs) across segments and axons. In contrast, in the axon with homogeneous myelination (Fig. 2I), segmental delays along the exposed parts remained constant. Taken together these insights hint at a non-linear relationship between the axonal conduction delay and the spatial heterogeneity of myelin sheathing.

### Heterogeneity of myelin sheathing impacts AP conduction reliability and susceptibility to failure

In the light of the above observations, a natural question arises as to how spatial heterogeneity of myelin sheaths influences the reliability, propagation and efficacy of AP conduction. The distinct fluctuations of AP waveform depicted in Fig. 2B-D suggest that the time required for an AP to traverse axons is likely contingent upon the precise arrangement and lengths of myelin sheaths and exposed segments. To quantify this path-dependence, we calculated the conduction delays and the corresponding CVs of axons with identical biophysical properties, for a fixed axonal length and for each value of PMC. The lengths and number of myelin sheaths along these axons were randomly sampled from either cortical or callosal myelination motifs (see Fig. 1).

As expected, axonal conduction delays decreased with the addition of myelin (Fig. 3A), while axonal CVs increased (Fig. 3B). While the mean axonal CVs and delays were found to be comparable between cortical and callosal myelination motifs (Fig. 3C), the dispersion in axonal conduction delays did exhibit important differences, indicative of variability in AP propagation between and even within cortical and callosal myelination motifs. To quantify this variability, we computed the dispersion (that is, standard deviation; std) of axonal conduction delays. We found that conduction delay variability was larger for callosal compared to cortical motifs, a trend that persisted across all PMC scenarios considered (Fig. 3D), despite being controlled for axonal length, PMC, and other physiological parameters (see Methods; Table 1). Furthermore, we observed a negative correlation between delay variability and PMC: as PMC increases, the dispersion of axonal conduction delays decreases. Another factor contributing to delay variability is the number of exposed segments: variability in axonal conduction delay is reduced whenever the number of exposed segments increases (Fig. 3F-G). Indeed, when the lengths of exposed segments decrease and become infinitesimal, as commonly assumed for nodes of Ranvier in many computational studies (***Fitzhugh, 1962; Goldman and Albus, 1968; Basser, 1993; Nygren and Halter, 1999; McIntyre et al., 2002; Gow and Devaux, 2008; Schmidt and Knösche, 2019; Naud and Longtin, 2019; Scurfield and Latimer, 2018; Babbs and Shi, 2013; Ashida and Nogueira, 2018; Grindrod and Sleeman, 1985***), conduction delay variability vanishes. Consistent with the findings above, cortical motifs demonstrated smaller variability compared to the corpus callosum. Interestingly, the associated axonal CVs also exhibited markedly lower variability in cortical compared to callosal motifs, whose CV was more variable across PMC values as shown in Fig. 3E. Taken together, these results confirm that regional differences in myelination significantly influence AP propagation variability.

**Figure 3.**
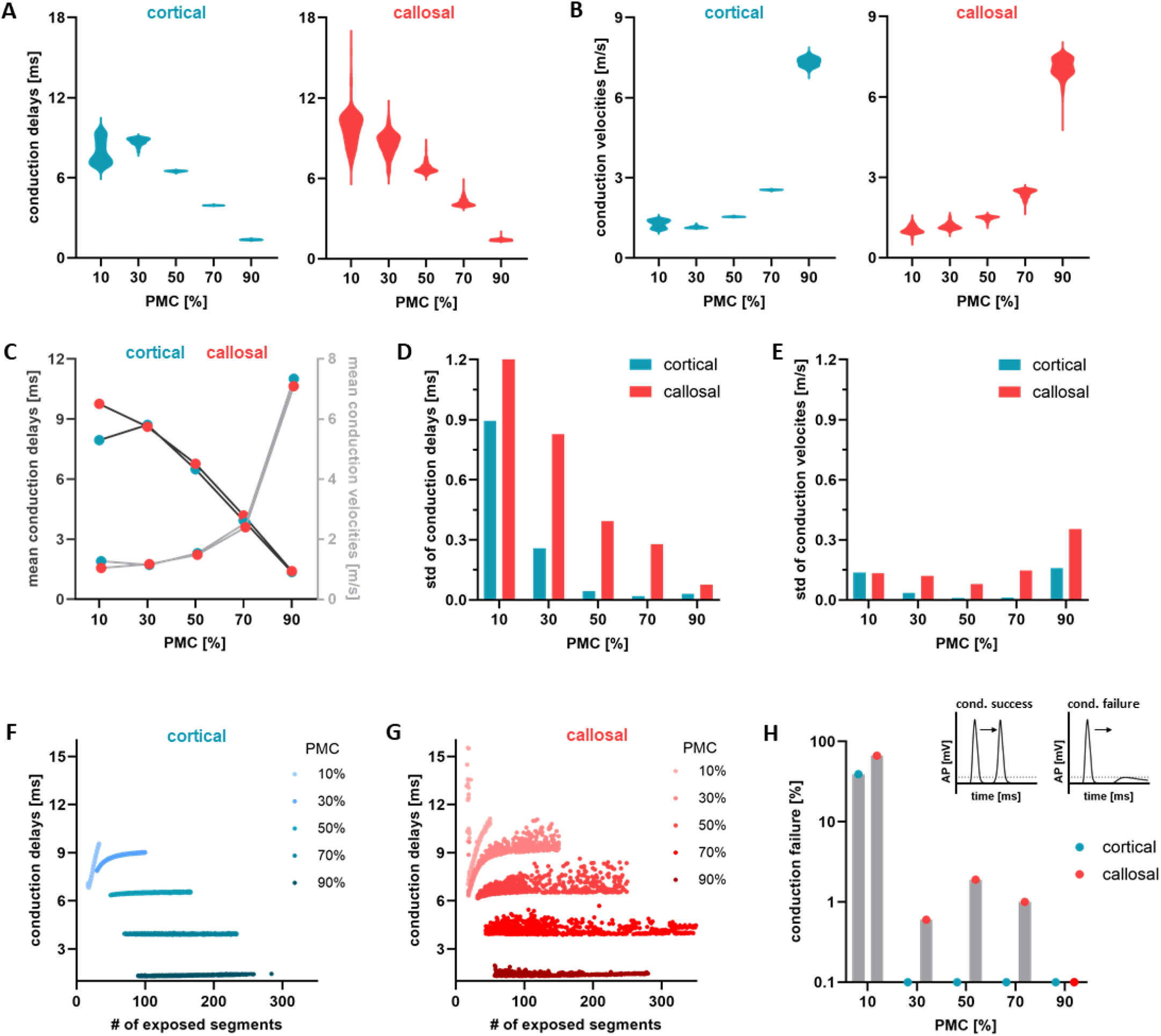
Axonal conduction variability as a result of spatially heterogeneous myelination along individual axons. **A**. Violin plots showing the distribution of axonal conduction delays (measured in *ms*) plotted for different myelin coverage marking the variability in axonal conduction caused by cortical and callosal myelin motifs. **B**. Violin plots of the corresponding distributions of axonal CVs (measured in *m/s*) for all five cases of PMC and for cortical and callosal myelin motifs. **C**. Mean values of axonal conduction delays (dark grey) and CVs (light gray) as a function of PMC. **D**. Bar plots of the standard deviation (std) of axonal conduction delays, for cortical and callosal myelin motifs, as a function of PMC. **E**. Bar plots of the std of axonal CVs as a function of PMC. **F-G**. Scatter plots of axonal conduction delays with respect to the number of exposed segments. In panel **F**, the different shades of blue point out the myelin intensity along axons with cortical myelin motifs, while in panel **G**, the different shades of red indicate myelin intensity along axons with callosal motifs. **H**. Rate of axonal conduction failures, that is, the number of failures over the total number of trials, for each PMC value. The top left intake plot is a representation of conduction success. The peak of the AP remains above the threshold at all times indicating that the AP reaches successfully the end-point of the axon. The top right intake plot is a representation of a conduction failure. The peak of the AP reaches a sub-threshold value making it unable to elicit a depolarization afterwards. The results shown are for *K* = 1000 independent axons.

The consequences of myelination motifs extend beyond issues of conduction time reliability. Our analysis reveals that the faithful transmission of APs (that is, conduction success or failure) is also highly dependent on the distribution of myelin. Conduction failure can arise due to factors such as high stimulation frequency and/or reduced capacity of membrane potentials to depolarize in regions deprived of myelin (***Schauf and Davis, 1974***). This has important functional implications, including the inability to elicit depolarization in postsynaptic cells. In our model, conduction failures occur when the AP fails to reach the paranodal junction while travelling along longer exposed segments (see Methods). As a result, the amplitude of the AP falls below the threshold necessary for spike initiation, as depicted in the embedded plot of Fig. 3H. An illustration of successful conduction is portrayed in Fig. 3H, where the AP amplitude maintains supra-threshold values throughout the entire axon. By calculating the rate of conduction failures (see Methods), we found that both cortical and callosal motifs exhibited systematic failure in cases of low myelination (PMC=10%, Fig. 3H). This phenomenon is primarily attributed to the presence of long exposed segments. These segments provide more opportunities for ion leakage across the axonal membrane, which, among other factors, contributes to the gradual decrease in depolarization amplitude necessary for AP propagation. A considerably lower rate of conduction failures was observed in axons exhibiting higher degrees of myelination, with myelination motifs inherent to the corpus callosum showing slightly greater susceptibility to failures. Such differences can be linked to the larger representation of long unmyelinated segments in callosal motifs (Fig. 1J). In summary, these findings suggest that failure susceptibility might increase as axons traverse the corpus callosum, while grey matter myelination motifs – inherently less heterogeneous – may demonstrate heightened resilience to changes in PMC.

### AP propagation is non-linear along heterogeneously myelinated axons

The results depicted in Figs. 2, 3 suggest that axonal conduction delays are not solely determined by PMC or total axonal length; rather, conduction timing depends on the specific placement, ordering, and lengths of myelin sheaths and exposed segments. Indeed, even when controlling for differences in PMC and axonal length, conduction delays demonstrate important variability directly attributable to cortical or callosal myelin sheaths arrangement (Fig. 3D). In Fig. 4A-C, we plotted segmental delays against the length of the corresponding segment. Given that APs propagate rapidly along myelin sheaths through diffusion (see Methods), such delays are predominantly due to propagation along exposed segments. For axons with homogeneous myelin sheathing, conduction time scales linearly with the lengths of exposed segments, except for very long segments (Fig. 4A). The dispersion of segmental delays is zero, and the CV, reflected in the slope of the curve, remains constant. In contrast, axons with cortical and callosal motifs show increased segmental delay dispersion proportional to myelin sheath heterogeneity and PMC. The CVs fluctuate across segments and axons, as evidenced by the important variability in the slopes of the curves in Fig. 4B-C. In line with our previous observations, callosal motifs did exhibit a more salient non-linearity combined with heightened variability of segmental delays, a feature directly linked to the statistical differences in exposed /myelinated segment lengths and number as well as the positioning of the segments on axons, compared to the cortical motifs (Fig. 1). These findings propose that axonal conduction is a path-dependent process, sensitive to cumulative transitions between myelinated and exposed segments of the axon, thereby confirming a dependence of AP propagation on myelination motifs.

**Figure 4.**
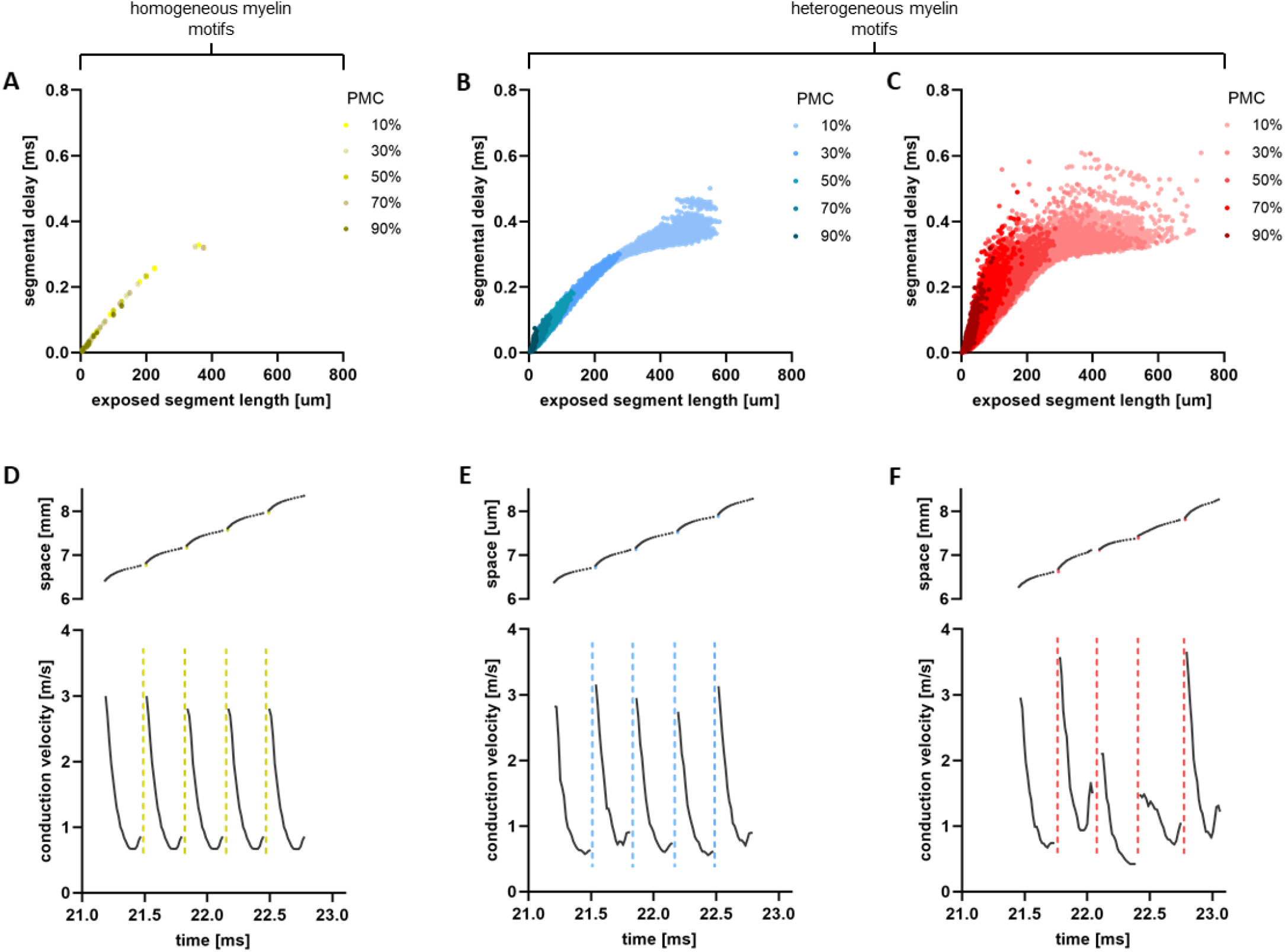
Non-linearity and path-dependence of AP propagation along axons with heterogeneous myelin sheathing. **A-C**. Conduction delays along individual exposed segments (that is, segmental delays) are plotted as a function of their individual lengths. In panel **A**, the model axons feature homogeneous myelin sheathing, while in panels **B** (blue; cortical) and **C** (red; callosal) myelin sheaths are spatially heterogeneous. In each panel, color shading indicates PMC: light colors refer to low PMC values, and darker colors indicate high PMC values. The plotted data are for *K* = 1000 independent axons for each of the myelination motifs and PMC. **D**. (top panel) For a section of an individual axon with homogeneous myelination, we plotted how the propagation time of AP changes in space (that is, along successive segments of the axon). The black dots represent the time within exposed segments and the yellow dots the time within myelin sheaths. (bottom panel) Corresponding CVs within each of the segments. The dashed yellow lines represent the asymptotic behaviour of the CVs within myelin sheaths. **E-F**. Similarly, for an axon with cortical (panel **E**) and callosal (panel **F**) myelination motif. For panels **D-F**, simulations are for individual axons of length *L* = 10mm, *P MC* = 10% and *N*_*my*_ = 25. Biophysical parameter values can be found in Table 1.

To better understand this non-linear behaviour, we examined in further detail AP propagation along individual segments (Fig. 4D-F). Along segments covered by myelin, transmission occurs instantaneously, as expected from saltatory conduction, while across exposed segments transmission slows down due to ionic currents (see Methods), leading to a non-linear increase of AP propagation time. In a homogeneously myelinated axon, AP propagation time follows a periodic pattern across segments, repeated throughout the entire axon (Fig. 4D – top panel). However, for heterogeneously myelinated axons, delays were found to increase irregularly along successive exposed segments, with more pronounced changes observed for the callosal motif, which we recall is more heterogeneous (Fig. 4E-F). These trends are further reflected in the corresponding changes in CV experienced by the AP as it propagates along individual segments. Along myelin sheaths, CV is extremely high, reflecting the instantaneous spike timing. In contrast, within exposed segments, CV rapidly decreases as the AP enters the segments, and slightly increases before reaching the junction of the next myelin sheath (Fig. 4D-F – bottom panels). Mirroring the trend seen for the conduction delay, CV follows a periodic pattern in the homogeneously myelinated axon (Fig. 4D). However, fluctuations in CV were found to be especially irregular for the callosal motif (Fig. 4F). Taken together, these results suggest that APs are subjected to bouts of deceleration and acceleration along exposed segments. These collectively result in a deeply non-linear, path-dependent process, by which AP conduction becomes highly sensitive to myelin sheathing heterogeneity. It is interesting to consider the potential implications of the observed non-linearity in conduction delays and velocities, for instance when myelination motifs undergo demyelination.

### Vulnerability of heterogeneous myelination motifs to demyelination

Upon the onset of demyelination, the integrity of myelin sheaths becomes compromised or completely breaks down, subsequently causing disruptions in the transmission of nerve impulses (***Schauf and Davis, 1974***). Such demyelination results from injury or loss of myelin sheaths and/or OLs, and occurs in numerous neurological (***Duncan and Radcliff, 2016***) and neuropsychiatric (***Takahashi et al., 2011; Fields, 2008***) disorders. Given the statistical differences in lengths, number and spatial distributions of myelin sheaths (Fig. 1), is AP conduction along axons with cortical or callosal motifs equally vulnerable to demyelination? To answer this question, we examined how cortical and callosal myelination motifs were impacted by myelin injury, and quantified their resulting predisposition to conduction failure. We modelled demyelination by systematically reducing the number of myelin sheaths while replacing those damaged segments with excitable ones exhibiting lower excitability compared to nodes of Ranvier. This was done to account for a pathological decrease in ion channel density (see Methods), and implemented by increasing the depolarization threshold along demyelinated segments. A schematic illustration portraying the demyelination process described above is presented in Fig. 5A. The reduction in the number of myelin sheaths naturally affects the PMC in the damaged axons. Specifically, we initially set the PMC for healthy axons at 90%, and 15% of the total number of myelin sheaths were successively removed in each iteration.

**Figure 5.**
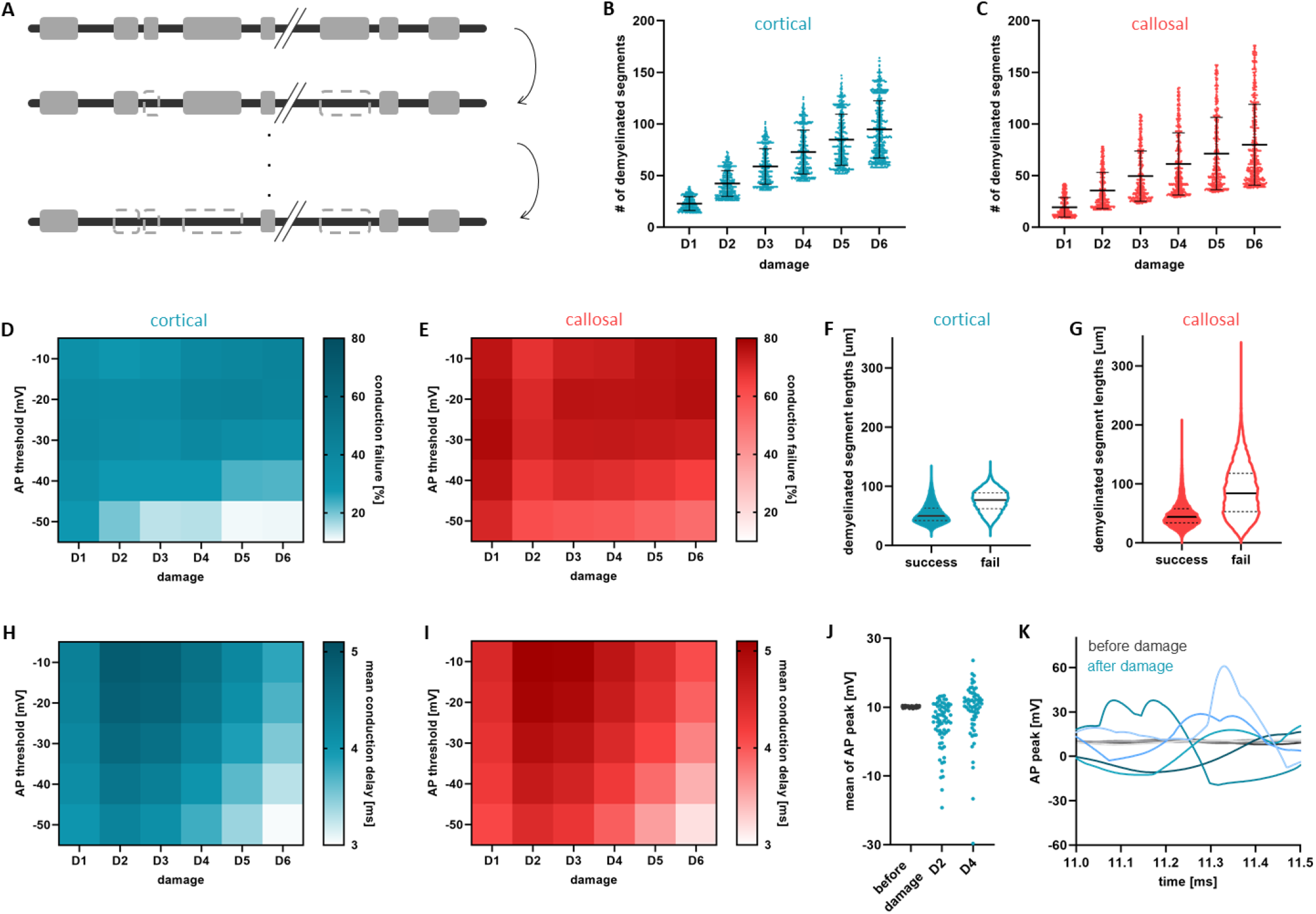
Vulnerability of cortical and callosal motifs to myelin injury. **A**. Schematic of the demyelination process. Axons of length *L* = 10mm and initial *P MC* = 90% undergo through successive myelin sheath removal, in which the number of myelin sheaths is reduced by 15% each time. This process is repeated six times to model different levels of severity of myelin damage (D1, …, D6). **B-C**. Scatter plots of the number of demyelinated segments along axons for all six steps of myelin damage for cortical (panel **B**) and callosal (panel **C**) myelination motifs. **D**. Heat map of the rate of conduction failure as a function of AP threshold and for each step of myelin damage for axons with cortical myelin motifs. **E**. Corresponding heat map of the rate of conduction failure for axons with callosal myelin motifs. **F-G**. Violin plots of demyelinated segment lengths characterizing AP conduction success and failure, for cortical and callosal motifs, respectively. **H-I**. Mean values of axonal conduction delays (only for those axons that successfully conduct) as a function of AP threshold and degree of myelin damage. Panel **H** is for axons featuring cortical motifs and panel **I** for callosal. **J**. Mean AP peak amplitude values for axons before damage and after damage (for steps D2 and D4 of demyelination) for axons with cortical myelination motifs in which successful conduction was observed. The plotted data for demyelinated axons are for AP threshold (only along the damaged segments) equal to −30mV. **K**. Representative examples of AP peak amplitude values for five axons before damage (grey shaded curves) and five axons after damage (blue shaded curves). All ten axons feature cortical motifs. Here damage is at D4 and AP threshold was set to −30mV.

Through this process, the number of demyelinated segments gradually increases (Fig. 5B-C). We note that on average, the number of demyelinated segments were roughly equal for both cortical and callosal myelin motifs.

To measure vulnerability, we computed the rate of axonal conduction failures for various AP threshold values and for successive application of myelin damage (Fig. 5D-E). We observed that callosal motifs were more vulnerable to demyelination, compared to cortical ones. This is confirmed by much higher rates of conduction failure (Fig. 5E). For both cortical and callosal motifs, conduction failures were expectedly found to be more prominent as the excitability of demyelinated segments decreases (that is, depolarization threshold increase, reflecting lower ion channel density). Closer examination of demyelinated sheath lengths in Fig. 5F-G indicates that conduction failures occur more frequently when damage/demyelination occurs along longer sheaths. In contrast, axons that succeed at conducting APs in spite of such damage are characterized by short demyelinated sheaths. We note that this is attributable to the fact that larger portions of the axon become exposed, and hence less excitable. Next, we examined how demyelination affected the conduction delays along axons that did not experience failure in spite of experiencing extensive damage. Figure 5H-I indicates an increase in mean conduction delay as the axon becomes increasingly demyelinated. This trend was equally observed in both cortical and callosal motifs. Intriguingly, our analysis predicts faster conduction with increased damage in some axons, suggesting a demyelination-induced increase in axonal CV. This counter-intuitive result could be attributed to the acceleration experienced by the AP as it propagates along exposed segments (Fig. 4D-F), as well as to other compensatory mechanisms pertaining to localized changes in axonal excitability (***Coggan et al., 2010***) (e.g., increase in effective excitability due to healthy neighboring segments/nodes with higher ion channel density). Nonetheless, these results suggest that in addition to failure, demyelination predisposes AP to time jitter by influencing their conduction time. To better understand this effect, we focused on axons exhibiting successful conduction (both before and after damage), and measured time fluctuations in the peak membrane potential (that is, states of maximal depolarization) as APs traverse the axons. As can be seen in Fig. 5J, APs were found to experience significant modulation when traversing damaged axons. The peak membrane potential varies considerably in damaged axons, while remains relatively constant throughout healthy axons (Fig. 5J-K). Such variability predisposes damaged axons to conduction block: demyelination-induced modulation in AP amplitude could fail to meet the depolarization threshold. As such, fluctuations in AP amplitude can be seen as interrogating both local excitability and myelin coverage, predisposing the axon to failure whenever the AP encounters long demyelinated segments (Fig. 5D-E). We highlight that this is another salient – and potentially pathological – manifestation of the path-dependent nature of axonal conduction along axons exhibiting heterogeneous myelin sheathing.

## Discussion

Numerous experimental studies have revealed that mammalian axons display a wide diversity of myelination patterns. Significant heterogeneity in the number, thickness, length, and distribution of myelin sheaths along and between different axons, as well as across brain regions (***Tomassy et al., 2014; Micheva et al., 2016; Hill et al., 2018; Call and Bergles, 2021; Auer et al., 2018; Stedehouder et al., 2019; Bakiri et al., 2011; Sturrock, 1980; Ford et al., 2015; Benamer et al., 2020; Chong et al., 2012***) has been observed. Notably, axons exhibit markedly different myelin sheathing motifs as they traverse the cortex and/or corpus callosum: cortical myelin sheaths display less variation compared to callosal ones, making them ideal candidates to examine the implications of myelin sheath heterogeneity on AP conduction (***Chong et al., 2012; Tomassy et al., 2014***). However, spatial heterogeneity in myelin sheathing is challenging to characterize with existing computational frameworks (***Fitzhugh, 1962; Goldman and Albus, 1968; Basser, 1993; Nygren and Halter, 1999; McIntyre et al., 2002; Gow and Devaux, 2008; Schmidt and Knösche, 2019; Naud and Longtin, 2019; Scurfield and Latimer, 2018; Babbs and Shi, 2013; Ashida and Nogueira, 2018; Grindrod and Sleeman, 1985***), often considering the axon as a cable with uniform myelin sheathing interspersed with exceedingly brief nodes of Ranvier. Motivated by the critical role played by myelin on AP conduction and its well-documented consequence on brain function (***Steadman et al., 2020; Pan et al., 2020; Bengtsson et al., 2005; Xin and Chan, 2020; McKenzie et al., 2014; Mabbott et al., 2006; Nagy et al., 2004; Pujol et al., 2006***), we here explored the influence of spatially heterogeneous myelin sheathing on axonal conduction reliability and predisposition to failure. Specifically, we developed a mathematical model of excitable axons displaying myelination motifs mirroring those observed experimentally in the cortex and corpus callosum of mice (***Chong et al., 2012***). We systematically examined how such myelination motifs influence AP conduction delays and CVs as well as vulnerability to failure in both healthy axons and those subjected to demyelination.

Our analysis revealed that variability in myelin patterns has an important impact on the reliability of axonal conduction. Indeed, axonal conduction was confirmed to be a nonlinear, path-dependent process, thereby sensitive to myelin sheaths arrangement through the cumulative effect of AP propagating across myelinated and/or exposed segments of various lengths. Variability in conduction delays was found to correlate inversely with myelin coverage and differed between callosal and cortical motifs. This variability was however found to be more prominent amongst callosal motifs. We emulated demyelinating damage by selectively removing sheaths and quantifying resulting changes in AP conduction. Callosal myelination motifs, in particular, were found to display a greater sensitivity to demyelination, exhibiting increased failure rates compared to cortical motifs.

The diversity in myelin sheaths distribution along axons is the manifestation of highly plastic and area-specific myelination processes, as well as the consolidation of signaling pathways by OLs. Through adaptive myelination and remodelling, myelin sheaths may retract and/or elongate (***Gibson et al., 2014; Baraban et al., 2018; Krasnow et al., 2018***), altering the lengths of exposed segments/nodes of Ranvier in an axon-specific manner (***Arancibia-Cárcamo et al., 2017; Koskinen et al., 2023***), thereby resulting in spatially heterogeneous myelination pattern. Myelin integrity has been linked to a wide array of brain functions ranging from memory (***Steadman et al., 2020; Pan et al., 2020***) and learning (***Bacmeister et al., 2022; Bengtsson et al., 2005; Xin and Chan, 2020; McKenzie et al., 2014***) to executive functions (***Mabbott et al., 2006; Nagy et al., 2004; Pujol et al., 2006***).

In contrast, compromised myelin integrity, resulting from the loss of myelin sheaths and/or Ols impairment, is linked to an increasing range of neurological (***Duncan and Radcliff, 2016; de Faria Jr et al., 2021***) and neuropsychiatric conditions (***Takahashi et al., 2011; Fields, 2008; de Faria Jr et al., 2021***). Aberrant and/or maladaptive changes such as the lengthening of the nodes of Ranvier, breakdown of the electrically insulating barrier between the myelin sheath and the axonal membrane at the paranodal region, and shrinkage of the sheaths leading to the exposure of K^+^ channels in the juctaparanodes, are also potential contributors to the modulation of myelinated axon functionality. These changes could result in a slowdown of the AP propagation or even a conduction failure across a diverse spectrum of pathological conditions (***Arancibia-Carcamo and Attwell, 2014; Dolma and Joshi, 2023; Freeman et al., 2016***).

Our results identify an important difference between myelination motifs in the cortex and corpus callosum, and their potential implications in myelin disorders. This difference echoes a long-standing conundrum in the literature pertaining to the pathological manifestation, consequences and clinical relevance of grey versus white matter lesions observed across multiple sclerosis (MS) stages (***Prins et al., 2015; Lie et al., 2022; Geurts and Barkhof, 2008***). It is interesting to conjecture that variations in myelination patterns and resulting differences in vulnerability to demyelinating damage reported here may be involved in MS progression. For instance, the relatively higher resilience of cortical myelination motifs to damage (cf. Fig. 5) allows us to hypothesize that grey matter could involved in later stages of the disease compared to the white matter. Additionally, recent findings have linked aberrant myelination of the corpus callosum (but not the cortex) and epilepsy progression in mice (***Knowles et al., 2022***). These results indicate that maladaptive myelination in the corpus callosum may predispose brain networks to seizures by amplifying pathological oscillations. Our predictions pertaining to the heightened sensitivity of callosal motifs to changes in myelin suggest that the corpus callosum might represent a network of high vulnerability to maladaptive changes in myelin. Further research is required to delineate the respective contributions of grey and white matter axons in severity and progression of myelin-related disorders.

While providing valuable insights, our study nonetheless faces limitations that should be acknowledged. First, we introduced variability exclusively to the longitudinal lengths of myelin sheaths and exposed segments, while keeping the axon radius constant and refraining from altering the myelin thickness. Introducing additional sources of variability, such as the g-ratio and/or axon diameter (***Talidou et al., 2021***) would provide a more comprehensive understanding of how these factors interact and influence axonal conduction, and could significantly expand the scope of our conclusions.

Second, we treated exposed segments of various lengths as nodes of Ranvier, assuming that ionic conductance does not change with length. Specifically, our model did not account for the different types of ion channels, their distribution along the exposed segments, or potential variation in their density within a given axon. These factors could affect the CV and variability of signal transmission. A detailed study incorporating ion channel dynamics and their spatial distribution could address this limitation.

Third, in characterizing the relationship between diverse myelination motifs and variability in axonal conduction delay, we used a phenomenological approach grounded in axonal geometry. This approach assumed infinite resistance along myelin sheaths and neglected the morphological structure of the paranodes. A more precise mathematical model, incorporating additional features at the paranodal junctions, would significantly enhance the physiological relevance of our model, and would allow us to better understand the role played by paradonal junctions in AP conduction, providing a more accurate representation of signal transmission.

## Methods and Materials

### Constructing axons with heterogeneous myelination motifs

We consider a cable of fixed length *L*, representing a bare axon. The cumulative length of the cable covered by myelin (*L*_*my*_) is determined by the formula *L*_*my*_ = *P MC ⋅ L*, where *P MC* represents the percentage of myelin coverage. The remaining part of the axon constitutes the exposed length (*L*_*exp*_), expressed as *L*_*exp*_ = *L* − *L*_*my*_. To construct a myelinated axon, we generate myelin sheaths, each of length 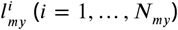, and exposed segments 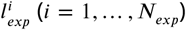, which are assembled in an alternating fashion. *N*_*my*_ and *N*_*exp*_ represent the total number of myelinated and exposed segments, respectively. Geometrically, the lengths *L*_*my*_ and *L*_*exp*_ correspond to the sum of individual myelinated or exposed segment lengths, as follows

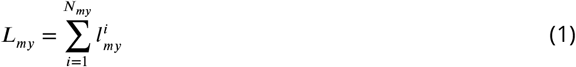

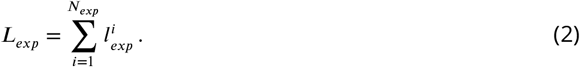

The distribution of myelin sheaths with variable lengths is carried out along the cable, as detailed in subsequent analyses.

#### Cortical myelination motifs

To create a cortical axon we generate a set of Poisson-distributed numbers that sum to *L*_*my*_. Each element of the set corresponds to a myelin sheath with the value of the element denoting the length of the sheath 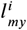. To generate this set, we first generate *L*_*my*_ points uniformly on the unit interval. In each sub-interval of length 1*/N*_*my*_, where *N*_*my*_ is the number of sheaths, there are on average *L*_*my*_*/N*_*my*_ events. Hence, there will be *N*_*my*_ events in total and the Poisson rate parameter becomes *λ* = *L*_*my*_*/N*_*my*_. We subsequently check whether the following constraints are satisfied: we require that the mean value of the sheath lengths is between 40 *μm* and 80 *μm* according to (***Chong et al., 2012; Call and Bergles, 2021***), and the minimal length is no smaller than 1 *μm* (***Call and Bergles, 2021***). The lengths of exposed segments were generated in a same manner, with the additional constraint that the minimum segment length is 1 *μm*, a value commonly associated with a node of Ranvier (***Arancibia-Cárcamo et al., 2017***).

#### Callosal myelination motifs

To create a callosal axon we adopt a similar strategy, but use a different distribution compared to the one used above. We generate a set of Gamma-distributed numbers (*N*_*my*_ in total) that sum to *L*_*my*_. The rationale underlying the selection of the Gamma distribution stems from the irregular sheath lengths generated per oligodendrocyte within the corpus callosum observed experimentally (***Chong et al., 2012***). Each element of a set following the Gamma distribution, Γ(*k, θ*), corresponds to a myelin sheath with the value of the element denoting the length of the sheath 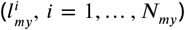. The shape parameter *k* and the scale parameter *θ* vary for each axon and are sampled from the uniform distribution. We have set two constraints for axons with callosal motifs: (i) the mean value of the sheath lengths is between 20 *μm* and 160 *μm*, and (ii) the minimal sheath length is greater or equal to 1 *μm*. We generate the exposed segment lengths in a same manner as with the myelin sheaths. The constraint is the same as for the cortical axons, namely that the minimal length of an exposed segment is 1 *μm*.

### Modelling AP propagation along heterogeneously myelinated axons

The membrane potential is determined by the properties and particular structure of myelinated and exposed segments along an axon. Due to high resistance along myelin sheaths, the membrane potential travels passively there and regenerates over exposed segments characterized by higher density of ion channels. Taking these properties into consideration, we built a model that governs the dynamics of the membrane potential along different segments of the axon. The model is defined by the following system of partial differential equations and conditions:

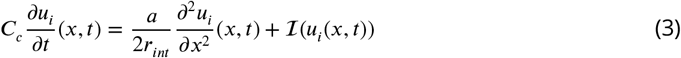

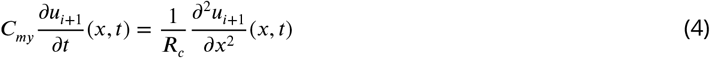

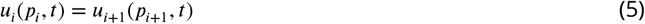

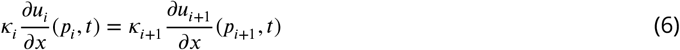

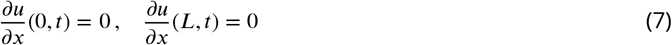

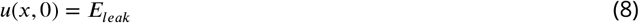

for *i* = 1, …, *N*, where *u*_*i*_(*x, t*), 0 < *x* < *L*, is the membrane potential within segment *i* (either myelinated or exposed/unmyelinated) at time *t* > 0 and for a total of *N* = *N*_*my*_ + *N*_*exp*_ segments.

The solution *u* of each differential equation involved in Eqs. (3)-(4) is the membrane potential within either an exposed segment/node of Ranvier (Eq. (3)) or a myelin sheath (Eq. (4)). We imposed continuity conditions at the paranodes (see Eqs. (5) - (6)), namely at the regions where the terminal myelin loops form septate-like junctions with the axolemma. Equation (5) indicates that the membrane potential *u* is continuous across a paranodal junction while Eq. (6) assumes continuity in flux, that is, the longitudinal currents should match at the junction. Following (***Rall, 1969***), we assume zero Neumann boundary conditions at either end (Eqs. (7)). For time *t* = 0 we set an initial condition equal to the leak reversal potential *E*_*leak*_ (Eq. (8)). We analyze the resulting governing equations and define all the parameters below.

#### Exposed segments/nodes of Ranvier

Equation (3) models the membrane potential *u* along the unmyelinated parts of the axon. This model has been introduced in Table 5 in (***Ashida and Nogueira, 2018***) as a simplified version of the Hodgkin-Huxley (***Hodgkin and Huxley, 1952***) and Wang-Buzaki models (***Wang and Buzsáki, 1996***). It accounts for spike generation and conduction, and has been inspired by the exponential integrate-and-fire model, which simulates the rapid growth of the membrane potential. In Eq. (3), the membrane capacitance of the cable is denoted by *C*_*c*_, *a* is the radius of the cable and *r*_*int*_ is the intracellular resistivity. Three currents are lumped together into ℐ(*u*): the leak current (*I*_*leak*_), the depolarizing current (*I*_*dep*_), and the repolarizing current (*I*_*rep*_). The leak current is given by

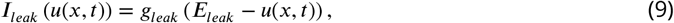

where *g*_*leak*_ is the leak conductance and *E*_*leak*_ the leak reversal potential. Both parameters were assumed to remain constant along the axon. The depolarizing current emulates the spike initiation and is defined as

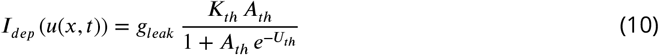

Where 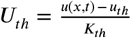. The parameter *u*_*th*_ is the spike threshold, and *K*_*th*_, *A*_*th*_ are parameters needed to control the exponential growth of the spike. When the membrane potential *u* reaches the preset value *u*_*rep*_, a repolarizing current is triggered forcing the membrane potential downward to the resting state. The repolarizing current is defined as:

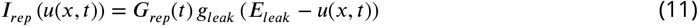

with *G*_*rep*_(*t*) being the repolarizing conductance for a time *t ≥ T*_*rep*_ (the time of activation of the repolarizing current):

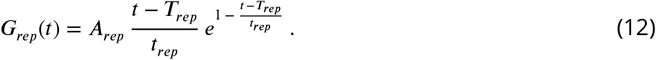

The parameters *A*_*rep*_ and *t*_*rep*_ are the amplitude and time constants of the repolarizing conductance, respectively.

To generate an AP, a brief external current *I*_*inj*_ is injected at a specific location at the beginning of the axon for a set duration *T*_*S*_:

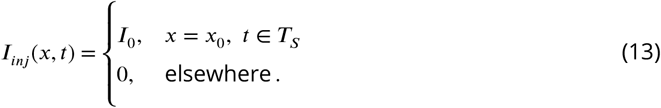

For a thorough discussion of the model used in Eq. (3), and its advantages and disadvantages compared to the classical conduction models, we refer the reader to the original work (***Ashida and Nogueira, 2018***). The parameter values used here are also taken from (***Ashida and Nogueira, 2018***) and are summarised in Table 1.

#### Myelin sheaths

Turning to the parts of the axon covered by myelin, we assume that the myelin sheaths have infinite resistance. Consequently, in the absence of ionic currents, the membrane potential *u* along a myelin sheath satisfies the diffusion equation Eq. (4) (cf. (***Dayan and Abbott, 2005***)). The capacitance of a myelin sheath *C*_*my*_ is defined as

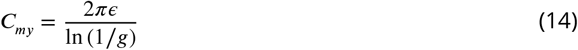

where *ϵ* is the permittivity of the myelin and *g* is the *g*-ratio, that is, the ratio of the inner-to-outer diameter of the myelinated axon. The cable resistance is given by the intracellular resistivity *r*_*int*_ over the cross-sectional area *πa*^2^, that is,

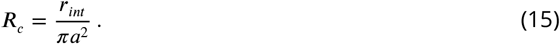

#### Paranodal junctions

As the myelin loops bind to the axonal membrane, they form paranodal axoglial junctions at both ends of every myelin sheath. While there is a rich selection of dynamics and functions at the paranodal junctions to maintain reliable saltatory conduction (***Rosenbluth, 2009***) we exclude those details from our model. We apply the continuity conditions given in Eqs. (5)-(6) at each paranodal junction, here denoted by *p*_*i*_. The diffusivities *κ* in Eq. (6) are defined by the diffusion coefficients of Eqs. (3)-(4). Specifically, from Eq. (3) we have 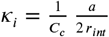 and from Eq. (4) 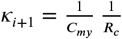.

### Axonal conduction delays, CVs and conduction failure

For each segment (either myelinated or exposed/unmyelinated) along an axon, we compute the difference between the time the membrane potential spikes at the beginning and end of the segment. This gives the conduction delay along individual segments referred thereafter as segmental delay. We denote these delays by 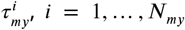 and 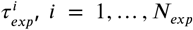 corresponding to the conduction times across individual myelinated and exposed segments, respectively. Then, the conduction delay *τ* along the entire axon is given by the sum of segmental delays:

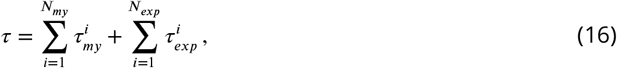

where we recall that *N*_*my*_ and *N*_*exp*_ denote the total number of myelinated and exposed segments along a single axon, respectively. Note that since APs propagate rapidly along myelin sheaths due to diffusion, the overall conduction delay *τ* in Eq. (16) predominantly reflects propagation time along exposed segments, that is,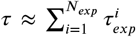. To obtain the corresponding conduction velocity (CV) we divide the total axonal length *L*, which is also equal to the sum of myelin sheaths and exposed segment lengths, by the conduction delay *τ*:

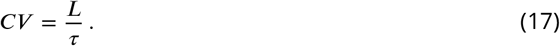

To examine whether an AP successfully conducts or fails to transverse the entire axon, we check the AP peak value at the end of each segment (myelinated or not). If the AP peak value is below the threshold value *u*_*th*_ (see Eq. (10)), then we have a conduction failure.

### Simulating myelin injury

We model demyelination of individual axons by reducing the number of myelin sheaths at a rate of 15%. Axons were demyelinated successively over a given number of discrete steps, repeated six times (D1, …, D6). In each step, 15% of the existing myelin sheaths were removed. Demyelination was conducted by randomly selecting myelin sheaths that had not been damaged in previous steps. Once a myelin sheath was damaged in a demyelination step, it remained impaired throughout subsequent iterations. In the event of myelin sheath damage, the damaged segment is replaced by an excitable one and modeled using Eq. (3) for exposed segments. Additionally, we vary the AP threshold value only at the injured segments to explore the implications of sodium channel density on axonal conduction. The AP threshold values range between −10 and −50 mV, all of which exceed the threshold at the unaffected exposed segments/nodes of Ranvier (−60.2 mV). The threshold value serves here as a proxy for the reduced excitability of the axon, reflecting a decrease in the density of sodium ion channels; an increase in channel density corresponds to a lower AP threshold.

We compute axonal conduction delays in a similar manner as in Eq. (16). Specifically, the conduction delay *τ* is determined by summing the conduction times across individual myelinated 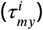, demyelinated 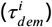 and exposed 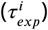 segments, so that

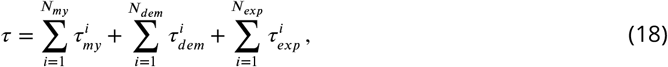

where *N*_*my*_, *N*_*dem*_ and *N*_*exp*_ denote the total number of myelinated, demyelinated and exposed segments, respectively.

### Numerical simulations

We modelled an axon as a medium composed of multiple layers, each representing the myelinated and exposed segments of the axon. Each segment was discretized into *n* + 1 equally spaced pieces, and second-order finite differences were used in space. Due to the varying lengths of the segments, the spatial step size *dx* differed for each segment. For temporal integration, we used the implicit Euler method.

To mitigate boundary effects at the beginning and end of the axon, we extend the axon at either end by creating copies of a portion of it with the same spatial properties. This approach allowed us to compute the conduction delays and velocities along fixed axonal lengths without boundary artifacts. The numerical schemes used for the approximation of the solution of Eqs. (3)-(8) were based on the methods proposed in (***Carr and Turner, 2016; Hickson et al., 2011***), which address the change of dynamics at the interfaces, namely at the paranodal junctions. The numerical scheme was implemented in a MATLAB code, which will be made available online upon publication.

## Acknowledgments

We thank the National Research Council of Canada (NSERC GRANT RGPIN-2017-06662) as well as the Canadian Institute of Health Research (CIHR GRANT NO PJT-156164) for funding.

## References

Arancibia-Carcamo IL, Attwell D. The node of Ranvier in CNS pathology. Acta Neuropathol. 2014; 128:161–175. https://link.springer.com/article/10.1007/s00401-014-1305-z, doi: 10.1007/s00401-014-1305-z.

Arancibia-Cárcamo IL, Ford MC, Cossell L, Ishida K, Tohyama K, Attwell D. Node of Ranvier length as a potential regulator of myelinated axon conduction speed. eLife. 2017 jan; 6:e23329. https://doi.org/10.7554/eLife.23329, doi: 10.7554/eLife.23329.

Ashida G, Nogueira W. Spike-Conducting Integrate-and-Fire Model. eNeuro. 2018; 5(4). https://www.eneuro.org/content/5/4/ENEURO.0112-18.2018, doi: 10.1523/ENEURO.0112-18.2018.

Auer F, Vagionitis S, Czopka T. Evidence for Myelin Sheath Remodeling in the CNS Revealed by In Vivo Imaging. Current Biology. 2018; 28(4):549–559.e3. https://www.sciencedirect.com/science/article/pii/S0960982218300198, doi: 10.1016/j.cub.2018.01.017.

Babbs CF, Shi R. Subtle paranodal injury slows impulse conduction in a mathematical model of myelinated axons. PLoS One. 2013; 8:e67767. https://journals.plos.org/plosone/article?id=10.1371/journal.pone.0067767, doi: 10.1371/journal.pone.0067767.

Bacmeister CM, Huang R, Osso LA, et al. Motor learning drives dynamic patterns of intermittent myelination on learning-activated axons. Nat Neurosci. 2022; 25:1300–1313. https://www.nature.com/articles/s41593-022-01169-4, doi: 10.1038/s41593-022-01169-4.

Bakiri Y, Káradóttir R, Cossell L, Attwell D. Morphological and electrical properties of oligodendrocytes in the white matter of the corpus callosum and cerebellum. The Journal of Physiology. 2011; 589(3):559–573. https://physoc.onlinelibrary.wiley.com/doi/abs/10.1113/jphysiol.2010.201376, doi: 10.1113/jphysiol.2010.201376.

Baraban M, Koudelka S, Lyons DA. Ca2+ activity signatures of myelin sheath formation and growth in vivo. Nat Neurosci. 2018; 21:19–23. https://doi.org/10.1038/s41593-017-0040-x, doi: 10.1038/s41593-017-0040-x.

Basser P. Cable equation for a myelinated axon derived from its microstructure. Med Biol Eng Comput. 1993; 31:S87–S92. https://link.springer.com/article/10.1007/BF02446655, doi: 10.1007/BF02446655.

Benamer N, Vidal M, Balia M, Angulo MC. Myelination of parvalbumin interneurons shapes the function of cortical sensory inhibitory circuits. Nat Commun. 2020; 11:5151. https://www.nature.com/articles/s41467-020-18984-7, doi: 10.1038/s41467-020-18984-7.

Bengtsson S, Nagy Z, Skare Sea. Extensive piano practicing has regionally specific effects on white matter development. Nat Neurosci. 2005; 8:1148–1150. https://www.nature.com/articles/nn1516, doi: 10.1038/nn1516.

Bradl M, Lassmann H. Oligodendrocytes: biology and pathology. Acta Neuropathol. 2010; 119:37–53. doi: 10.1007/s00401-009-0601-5.

Call CL, Bergles DE. Cortical neurons exhibit diverse myelination patterns that scale between mouse brain regions and regenerate after demyelination. Nat Commun. 2021; 12. https://www.nature.com/articles/s41467-021-25035-2, doi: 10.1038/s41467-021-25035-2.

Carr EJ, Turner IW. A semi-analytical solution for multilayer diffusion in a composite medium consisting of a large number of layers. Applied Mathematical Modelling. 2016; 40(15):7034–7050. https://www.sciencedirect.com/science/article/pii/S0307904X16301160, doi: 10.1016/j.apm.2016.02.041.

Chong SYC, Rosenberg SS, Fancy SPJ, Zhao C, Shen YAA, Hahn AT, McGee AW, Xu X, Zheng B, Zhang LI, Rowitch DH, Franklin RJM, Lu QR, Chan JR. Neurite outgrowth inhibitor Nogo-A establishes spatial segregation and extent of oligodendrocyte myelination. Proceedings of the National Academy of Sciences. 2012; 109(4):1299–1304. https://www.pnas.org/doi/abs/10.1073/pnas.1113540109, doi: 10.1073/pnas.1113540109.

Coggan JS, Prescott SA, Bartol TM, Sejnowski TJ. Imbalance of ionic conductances contributes to diverse symptoms of demyelination. Proceedings of the National Academy of Sciences. 2010; 107(48):20602–20609. https://www.pnas.org/doi/abs/10.1073/pnas.1013798107, doi: 10.1073/pnas.1013798107.

Dayan P, Abbott LF. Theoretical Neuroscience. MIT Press, Cambdridge. 2005; https://mitpress.mit.edu/9780262041997/theoretical-neuroscience/.

Dolma S, Joshi A. The Node of Ranvier as an Interface for Axo-Glial Interactions: Perturbation of Axo-Glial Interactions in Various Neurological Disorders. J Neuroimmune Pharmacol. 2023; 18:215–234. https://link.springer.com/article/10.1007/s11481-023-10072-z, doi: 10.1007/s11481-023-10072-z.

Duncan ID, Radcliff AB. Inherited and acquired disorders of myelin: The underlying myelin pathology. Experimental Neurology. 2016; 283:452–475. https://www.sciencedirect.com/science/article/pii/S0014488616300838, doi: 10.1016/j.expneurol.2016.04.002.

de Faria Jr O, Pivonkova H, Varga B, Timmler S, Evans KA, Káradóttir RT. Periods of synchronized myelin changes shape brain function and plasticity. Nature Neuroscience. 2021; 24(11):1508–1521.

Fields RD. White matter in learning, cognition and psychiatric disorders. Trends in Neuro-sciences. 2008; 31(7):361–370. https://www.sciencedirect.com/science/article/pii/S016622360800132X, doi: 10.1016/j.tins.2008.04.001.

Fitzhugh R. Computation of Impulse Initiation and Saltatory Conduction in a Myelinated Nerve Fiber. Biophysical Journal. 1962; 2(1):11–21. https://www.sciencedirect.com/science/article/pii/S0006349562868374, doi: 10.1016/S0006-3495(62)86837-4.

Ford MC, Alexandrova O, Cossell L, Stange-Marten A, Sinclair J, Kopp-Scheinpflug M C Pecka, Attwell D, Grothe B. Tuning of Ranvier node and internode properties in myelinated axons to adjust action potential timing. Nat Commun. 2015; 6:8073. https://www.nature.com/articles/ncomms9073, doi: 10.1038/ncomms9073.

Freeman SA, Desmazières A, Fricker D, Lubetzki C, Sol-Foulon N. Mechanisms of sodium channel clustering and its influence on axonal impulse conduction. Cell Mol Life Sci. 2016; 73:723–735. https://link.springer.com/article/10.1007/s00018-015-2081-1, doi: 10.1007/s00018-015-2081-1.

Geurts JJ, Barkhof F. Grey matter pathology in multiple sclerosis. The Lancet Neurology. 2008; 7(9):841–851. https://www.sciencedirect.com/science/article/pii/S1474442208701911, doi: 10.1016/S1474-4422(08)70191-1.

Gibson EM, Purger D, Mount CW, Goldstein AK, Lin GL, Wood LS, Inema I, Miller SE, Bieri G, Zuchero JB, Barres BA, Woo PJ, Vogel H, Monje M. Neuronal Activity Promotes Oligodendrogenesis and Adaptive Myelination in the Mammalian Brain. Science. 2014; 344(6183):1252304. https://www.science.org/doi/abs/10.1126/science.1252304, doi: 10.1126/science.1252304.

Goldman L, Albus JS. Computation of Impulse Conduction in Myelinated Fibers; Theoretical Basis of the Velocity-Diameter Relation. Biophysical Journal. 1968; 8(5):596–607. https://www.sciencedirect.com/science/article/pii/S0006349568865105, doi: 10.1016/S0006-3495(68)86510-5.

Gow A, Devaux J. A model of tight junction function in central nervous system myelinated axons. Neuron Glia Biol. 2008; 4:307–317. https://www.cambridge.org/core/journals/neuron-glia-biology/article/model-of-tight-junction-function-in-central-nervous-system-myelinated-axons/A13094DCBE92B9720396656B82B1E46B, doi: 10.1017/S1740925X09990391.

Grindrod P, Sleeman BD. A model of a myelinated nerve axon: threshold behaviour and propagation. J Math Biol. 1985; 23:119–135. https://link.springer.com/article/10.1007/BF00276561, doi: 10.1007/BF00276561.

Hickson RI, Barry SI, Mercer GN, Sidhu HS. Finite difference schemes for multilayer diffusion. Mathematical and Computer Modelling. 2011; 54(1):210–220. https://www.sciencedirect.com/science/article/pii/S0895717711000938, doi: 10.1016/j.mcm.2011.02.003.

Hill RA, Li AM, Grutzendler J. Lifelong cortical myelin plasticity and age-related degeneration in the live mammalian brain. Nat Neurosci. 2018; 21:683–695. https://www.nature.com/articles/s41593-018-0120-6, doi: 10.1038/s41593-018-0120-6.

Hodgkin AL, Huxley AF. A quantitative description of membrane current and its application to conduction and excitation in nerve. The Journal of Physiology. 1952; 117(4):500–544. https://physoc.onlinelibrary.wiley.com/doi/abs/10.1113/jphysiol.1952.sp004764, doi: 10.1113/jphysiol.1952.sp004764.

Knowles JK, Xu H, Soane C, et al. Maladaptive myelination promotes generalized epilepsy progression. Nat Neurosci. 2022; 25:596–606. https://doi.org/10.1038/s41593-022-01052-2, doi: 10.1038/s41593-022-01052-2.

Koskinen MK, Laine M, Abdollahzadeh A, Gigliotta A, Mazzini G, Journée S, Alenius V, Trontti K, Tohka J, Hyytiä P, Sierra A I H. Node of Ranvier remodeling in chronic psychosocial stress and anxiety. Neuropsychopharmacology. 2023; 48:1532–1540. https://www.nature.com/articles/s41386-023-01568-6, doi: 10.1038/s41386-023-01568-6.

Krasnow AM, Ford MC, Valdivia LE, Wilson SW, Attwell D. Regulation of developing myelin sheath elongation by oligodendrocyte calcium transients in vivo. Nat Neurosci. 2018; 21:24–28. https://www.nature.com/articles/s41593-017-0031-y, doi: 10.1038/s41593-017-0031-y.

Lie IA, Weeda MM, Mattiesing RM, Mol MAE, Pouwels PJW, Barkhof F, Øivind Torkildsen, Bø L, Myhr KM, Vrenken H. Relationship Between White Matter Lesions and Gray Matter Atrophy in Multiple Sclerosis. Neurology. 2022; 98(15):e1562–e1573. https://www.neurology.org/doi/abs/10.1212/WNL.0000000000200006, doi: 10.1212/WNL.0000000000200006.

Mabbott DJ, Noseworthy M, Bouffet E, Laughlin S, Rockel C. White matter growth as a mechanism of cognitive development in children. NeuroImage. 2006; 33(3):936–946. https://www.sciencedirect.com/science/article/pii/S1053811906007865, doi: 10.1016/j.neuroimage.2006.07.024.

McIntyre CC, Richardson AG, Grill WM. Modeling the Excitability of Mammalian Nerve Fibers: Influence of Afterpotentials on the Recovery Cycle. Journal of Neurophysiology. 2002; 87(2):995–1006. https://doi.org/10.1152/jn.00353.2001, doi: 10.1152/jn.00353.2001.

McKenzie IA, Ohayon D, Li H, de Faria JP, Emery B, Tohyama K, Richardson WD. Motor skill learning requires active central myelination. Science. 2014; 346:318–322. https://www.science.org/doi/10.1126/science.1254960, doi: 10.1126/science.1254960.

Micheva KD, Wolman D, Mensh BD, Pax E, Buchanan J, Smith SJ, Bock DD. A large fraction of neocortical myelin ensheathes axons of local inhibitory neurons. eLife. 2016 jul; 5:e15784. https://doi.org/10.7554/eLife.15784, doi: 10.7554/eLife.15784.

Nagy N, Westerberg H, Klingberg T. Maturation of white matter is associated with the development of cognitive functions during childhood. J Cogn Neurosci. 2004; 16:1227–1233. https://direct.mit.edu/jocn/article/16/7/1227/3907/Maturation-of-White-Matter-is-Associated-with-the, doi: 10.1162/0898929041920441.

Naud R, Longtin A. Linking demyelination to compound action potential dispersion with a spike-diffuse-spike approach. J Math Neurosci. 2019; 9. https://mathematical-neuroscience.springeropen.com/articles/10.1186/s13408-019-0071-6, doi: 10.1186/s13408-019-0071-6.

Nygren A, Halter JA. A General Approach to Modeling Conduction and Concentration Dynamics in Excitable Cells of Concentric Cylindrical Geometry. Journal of Theoretical Biology. 1999; 199(3):329–358. https://www.sciencedirect.com/science/article/pii/S0022519399909621, doi: 10.1006/jtbi.1999.0962.

Pan S, Mayoral SR, Choi HS, Chan JR, Kheirbek MA. Preservation of a remote fear memory requires new myelin formation. Nat Neurosci. 2020; 23:487–499. https://www.nature.com/articles/s41593-019-0582-1, doi: 10.1038/s41593-019-0582-1.

Prins M, Schul E, Geurts J, van der Valk P, Drukarch B, van Dam AM. Pathological differences between white and grey matter multiple sclerosis lesions. Annals of the New York Academy of Sciences. 2015; 1351(1):99–113. https://nyaspubs.onlinelibrary.wiley.com/doi/abs/10.1111/nyas.12841, doi: 10.1111/nyas.12841.

Pujol J, Soriano-Mas C, Ortiz H, Sebastián-Gallés N, Losilla JM, Deus J. Myelination of language-related areas in the developing brain. Neurology. 2006; 66(3):339–343. https://www.neurology.org/doi/abs/10.1212/01.wnl.0000201049.66073.8d, doi: 10.1212/01.wnl.0000201049.66073.8d.

Rall W. Time Constants and Electrotonic Length of Membrane Cylinders and Neurons. Biophysical Journal. 1969; 9(12):1483–1508. https://www.sciencedirect.com/science/article/pii/S0006349569864672, doi: 10.1016/S0006-3495(69)86467-2.

Rosenbluth J. Multiple functions of the paranodal junction of myelinated nerve fibers. Journal of Neuroscience Research. 2009; 87(15):3250–3258. https://onlinelibrary.wiley.com/doi/abs/10.1002/jnr.22013, doi: 10.1002/jnr.22013.

Schauf CL, Davis FA. Impulse conduction in multiple sclerosis: a theoretical basis for modification by temperature and pharmacological agents. Journal of Neurology, Neurosurgery & Psychiatry. 1974; 37(2):152–161. https://jnnp.bmj.com/content/37/2/152, doi: 10.1136/jnnp.37.2.152.

Schmidt H, Knösche TR. Action potential propagation and synchronisation in myelinated axons. PLoS Comput Biol. 2019; 15:e1007004. https://journals.plos.org/ploscompbiol/article?id=10.1371/journal.pcbi.1007004, doi: 10.1371/journal.pcbi.1007004.

Scurfield A, Latimer DC. A computational study of the impact of inhomogeneous internodal lengths on conduction velocity in myelinated neurons. PLoS One. 2018; 13:e0191106. https://journals.plos.org/plosone/article?id=10.1371/journal.pone.0191106, doi: 10.1371/journal.pone.0191106.

Simons M, Nave KA. Oligodendrocytes: Myelination and Axonal Support. Cold Spring Harb Perspect Biol. 2015; 22:a020479. doi: 10.1101/cshperspect.a020479.

Steadman PE, Xia F, Ahmed M, Mocle AJ, Penning ARA, Geraghty AC, Steenland HW, Monje M, Josselyn SA, Frankland PW. Disruption of Oligodendrogenesis Impairs Memory Consolidation in Adult Mice. Neuron. 2020; 105(1):150–164.e6. https://www.sciencedirect.com/science/article/pii/S0896627319308864, doi: 10.1016/j.neuron.2019.10.013.

Stedehouder J, Brizee D, Slotman JA, Pascual-Garcia M, Leyrer ML, Bouwen BL, Dirven CM, Gao Z, Berson DM, Houtsmuller AB, Kushner SA. Local axonal morphology guides the topography of interneuron myelination in mouse and human neocortex. eLife. 2019 nov; 8:e48615. https://doi.org/10.7554/eLife.48615, doi: 10.7554/eLife.48615.

Sturrock RR. Myelination of the mouse corpus callosum. Neuropathology and Applied Neurobiology. 1980; 6(6):415–420. https://onlinelibrary.wiley.com/doi/abs/10.1111/j.1365-2990.1980.tb00219.x, doi: 10.1111/j.1365-2990.1980.tb00219.x.

Takahashi N, Sakurai T, Davis KL, Buxbaum JD. Linking oligodendrocyte and myelin dysfunction to neurocircuitry abnormalities in schizophrenia. Progress in Neurobiology. 2011; 93(1):13–24. https://www.sciencedirect.com/science/article/pii/S0301008210001693, doi: 10.1016/j.pneurobio.2010.09.004.

Talidou A, Burchard A, Sigal IM. Near-Pulse Solutions of the FitzHugh-Nagumo Equations on Cylindrical Surfaces. Journal of NonLinear Science. 2021; 31(3):57. https://link.springer.com/article/10.1007/s00332-021-09710-8, doi: 10.1007/s00332-021-09710-8.

Talidou A, Frankland PW, Mabbott D, Lefebvre J. Homeostatic coordination and up-regulation of neural activity by activity-dependent myelination. Nat Comput Sci. 2022; 2:665–676. https://www.nature.com/articles/s43588-022-00315-z, doi: 10.1038/s43588-022-00315-z.

Tomassy GS, Berger DR, Chen HH, Kasthuri N, Hayworth KJ, Vercelli A, Seung HS, Lichtman JW, Arlotta P. Distinct Profiles of Myelin Distribution Along Single Axons of Pyramidal Neurons in the Neocortex. Science. 2014; 344(6181):319–324. https://www.science.org/doi/abs/10.1126/science.1249766, doi: 10.1126/science.1249766.

Wang XJ, Buzsáki G. Gamma Oscillation by Synaptic Inhibition in a Hippocampal Interneuronal Network Model. Journal of Neuroscience. 1996; 16(20):6402–6413. https://www.jneurosci.org/content/16/20/6402, doi: 10.1523/JNEUROSCI.16-20-06402.1996.

Xin W, Chan JR. Myelin plasticity: sculpting circuits in learning and memory. Nat Rev Neurosci. 2020; 21:682–694. https://www.nature.com/articles/s41583-020-00379-8, doi: 10.1038/s41583-020-00379-8.

